## Supplementary Material for "Emergence of Supercoiling-Mediated Regulatory Networks through the Evolution of Bacterial Chromosome Organization"

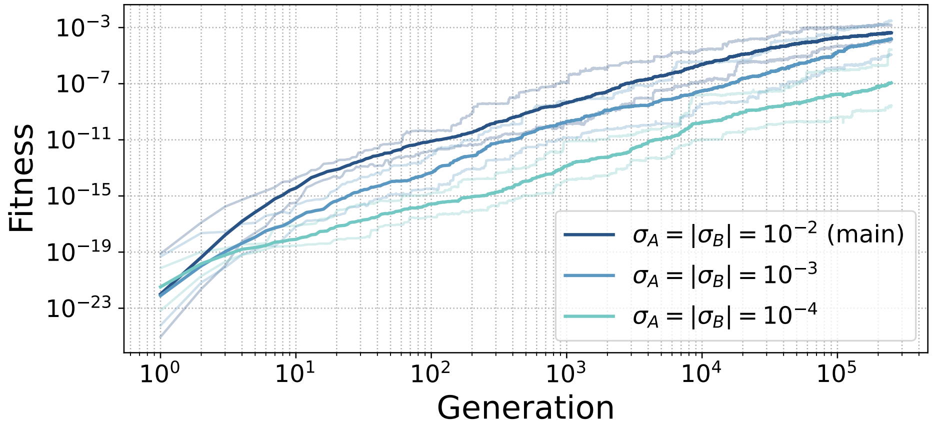


Figure S1: Average fitness during evolution, with environmental shifts in supercoiling logarithmically decreasing in absolute value: $\sigma_A = 10^{-2}$ and $\sigma_B = -10^{-2}$ (main run), $\sigma_A = 10^{-3}$ and $\sigma_B = -10^{-3}$ (10 times smaller than in the main run) and $\sigma_A = 10^{-4}$ and $\sigma_B = -10^{-4}$ (100 times smaller than the main run). Lighter lines represent the first and last decile of the data.

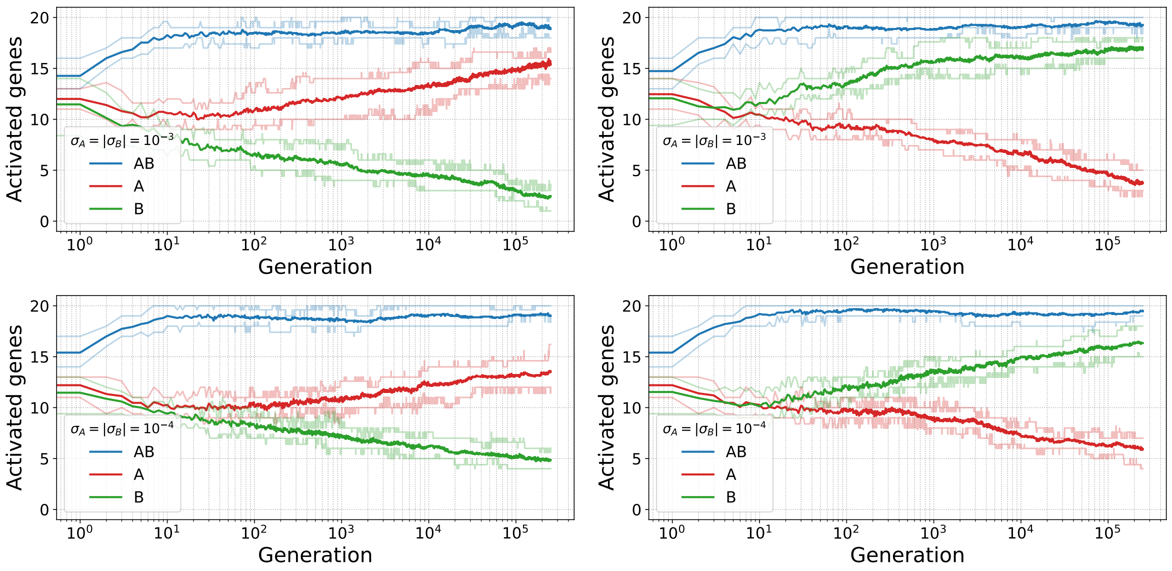


Figure S2: Evolution of the number of activated genes in environment A (left) and environment B (right), with environmental supercoiling shifts $\sigma_A = 10^{-3}$ and $\sigma_B = -10^{-3}$ (top) and $\sigma_A = 10^{-4}$ and $\sigma_B = -10^{-4}$ (bottom). Lighter lines represent the first and last decile of the data.

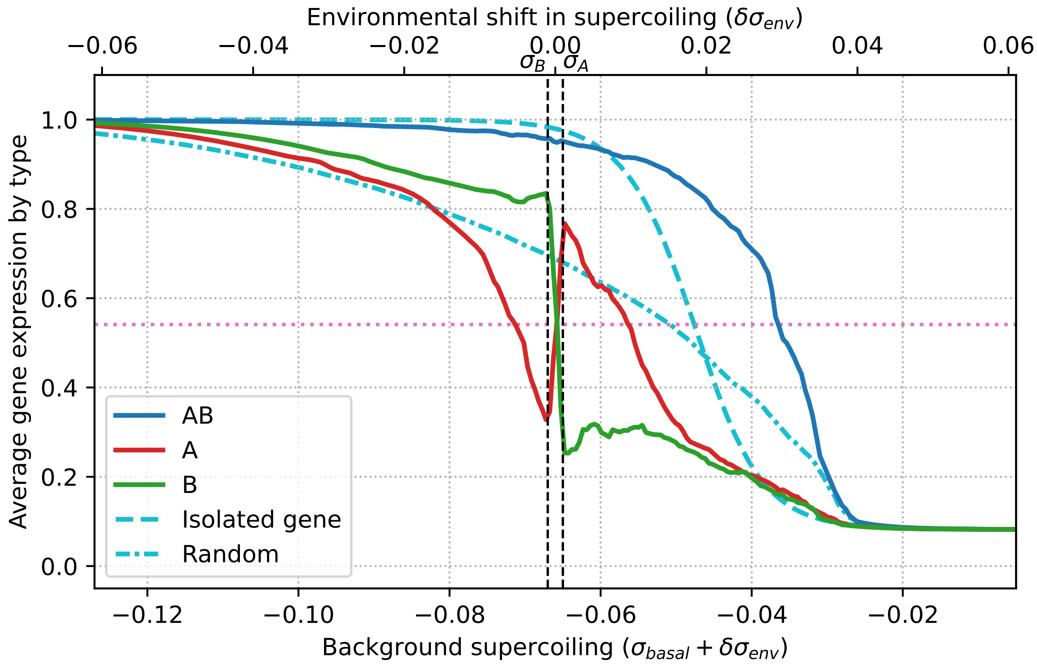


Figure S3: Average gene expression as a function of background supercoiling, with environmental supercoiling shifts $\sigma_A = 10^{-3}$ and $\sigma_B = -10^{-3}$.

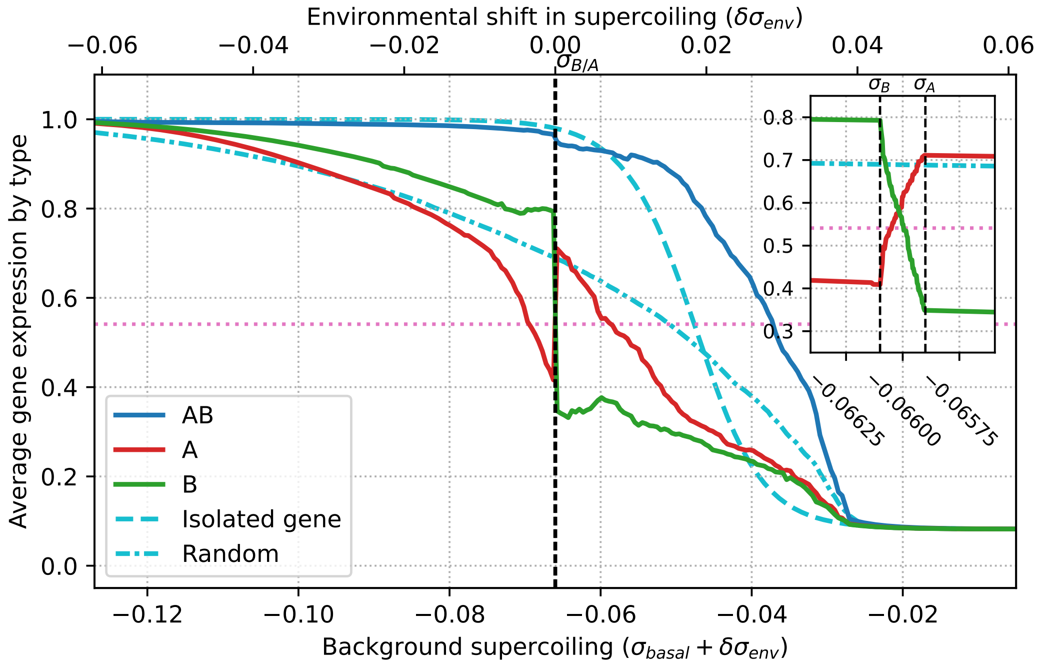


Figure S4: Average gene expression as a function of background supercoiling, with environmental supercoiling shifts $\sigma_A = 10^{-4}$ and $\sigma_B = -10^{-4}$. The inset at the top right of the figure shows a 150x zoom on supercoiling shift values near zero.
